# Optimizing multi environment cowpea yield trial by appraising the fitness and similarities among test sites: a case study of IITA cowpea breeding program

**DOI:** 10.64898/2026.01.21.700123

**Authors:** Oyebode Gideon Oluwaseye, ThankGod Oche Ogwuche

**Affiliations:** International Institute of Tropical Agriculture (IITA), IITA Kano Station, PMB 3112, Kano, Nigeria; International Institute of Tropical Agriculture (IITA), Ibadan Station PMB 5320, Ibadan, Nigeria

**Keywords:** GGE-Biplot, Best linear unbiased predictor (BLUP), Discriminativeness and representativeness, Multi-environment trial, Testing efficiency, Testing networks, Linear mixed model, G x E, Test site similarity

## Abstract

Optimization of Multi environment trials (MET) trials require an efficient testing network consisting of ideal sites which should be highly discriminative and representative of their mega environments. Also concurrently managing sites that are similar in terms of information there give about genotype performance adds no extra utility to the breeding program. The objective of this study was to appraise the testing network of the cowpea breeding program of the International Institute of tropical Agriculture (IITA) based in Nigeria on the basis of their representativeness and discriminatory ability for grain yield, investigate the similarities among them and assess the variance components. 6 set sets of Advance yield trial with unique entries were analysed using Mixed models, best linear unbiased estimate (BLUPs) and BLUP based GGE biplot. Results showed significant means squares Genotype x Location interaction effect for all 6 sets; this justifies the to study GEI in cowpea, partitioning variance showed that Environment main effect accounted for the largest proportion of the total phenotypic variance, with a range of 58.1 (Adv 4) to 73.9% (Adv 3), Environment was followed by Genotype x Location interaction which explained between 16.1% (Adv 5) to 22.6% (Adv 4). Genotype main effect accounted for the least with a range of 9.5% (Adv 6) to 19.3% (Adv 4). Broad sense-heritability for GY was high in Advance 1-5 (ranging from 0.62-0.70) and medium in advance 6 (0.53). Shika June, Shika August and Minjibir consistently showed high discriminativeness with Minjibir been the most discriminating environment for this study. Ibadan September and Ibadan May on the other hand consistently showed poor discriminating ability while BUK was inconsistent having good discriminative ability in Sets 1 and 3 while it was poor in sets 2, 5 and 6. In terms of representativeness, no environment consistently had desirable results however, Shika August and Shika June were most representative while Minjibir was least representative. On the basis of discriminativeness and representativeness, the six environments were ranked in other of desirability as Shika August> Shika June> Minjibir > Ibadan September > BUK farm > Ibadan May. In terms of similarity among environments, both the BLUP-Based GGE biplot and genotypic correlations indicated that Ibadan May and September were consistently grouped together and highly correlated; indicating that they are similar, while, Shika August and Shika June were found to be unique. Ibadan May was therefore adjudged a redundant location and could be dropped and replaced without any loss of accuracy because it was neither discriminative nor representative in all biplots draw and it consistently fell in the same Mega environment with Ibadan September. We concluded that there is a need to sample more testing sites and validate their fitness for multi environment yield trials using methods applied in this study.

## Introduction

The present epoch of climate change aggravated by global warming places a strain on the effort to attain food and nutrition security; especially in Sub-Saharan Africa (SSA). Cowpea (Vigna unguiculata L. Walp.) (2n = 22) is the most important leguminous crop for promoting food and nutrition security in SSA (Ishikawa et al., 2020, Boukar et al. 2011, Fatokun et al. 2002) where it is already the main source of plant-based protein (Muranaka et al., 2015) and possesses exceptional adaptation and tolerance to environmental stresses unique to SSA (Murdock et al., 2008). Being a multi-purpose crop, it delivers excellent fodder for livestock of smallholder farmers with little inputs (Alemu et al., 2016) and fixes atmospheric nitrogen via root nodule rhizobia symbiosis, playing a crucial role in soil fertility and amendment. It is an important source of income to farmers in SSA; where it is grown as a cash crop (Kormawa et al., 2002). Globally, cowpea is being cultivated on about 14.4 million hectares, with a world production of 8.9 million tons (MT) and Africa accounted for 95.2% (FAO STAT 2019). Nigeria is the largest producer of the crop with 2.7 MT, followed by Niger with 1.06MT (FAO stat, 2020).

Remarkable yield gap has been reported by recent surveys conducted in SSA (Erana et al., 2020, Mohamed et al., 2021, Bolarinwa et al 2021, Anago et al.,2021, Kouyate et al 2021), indicating that the productivity of the crop on farmer’s field is very low. This low productivity has been attributed to lack of improved and well adapted varieties that suits the farmers’ socioeconomic and environmental needs (Edward et al., 2021, Mohamed et al., 2021). The process of identifying outstanding varieties that suits the farmers’ needs is always disturbed by the ever-present Genotype by environment interaction (GEI) which causes inconsistent genotype performance with change in environment; complicating identification and recommendation of superior genotypes. This justifies the need for multi environment trials (METs) across diverse location and seasons (years) to ensure that only high yielding and stable genotypes are identified and released. The efficiency of METs depends on the suitability of the test environments for genotype evaluation. A good environment for METs must be discriminating of genotypes and representative of a growing region (Yan et al., 2007). Hence, plant breeding programs are required to frequently appraise the suitability of their test sites to ensure that no site is redundant and that each test site provides unique information about genotype performance. Also important, is the need to have proper information on the sensitivity, magnitude and type of GEI in play for grain yield as this will aid in implementing better breeding strategy (Ongom et al., 2021, Falcon et al., 2020, Badu-Apraku et at., 2011).

Prediction of genotype performance in a MET based on single-year multiple location data have been demonstrated to be as accurate as multiple years (Yan and Rajcan 2003, Cross and Helm 1986, Gellner 1989, and Bowman, 1998). Also, the utilization of Best linear unbiased estimates (BLUPs) further strengthens the accuracy that is obtained from single-year multiple location data. This is possible because the Mixed models permits the use of unbalanced data (typical of must METs) and can allow the estimation of BLUP (Yan and Rajcan 2003, Yan et al., 2002 ), in this study we explored the application of BLUPs to obtain GGE biplots. This was desirable approach, considering that the BLUP has the property of attenuation toward the grand mean, it made more statistical sense to apply it to MET data sets that were unbalanced (Piepho et al., 2008, Piepho, 1994; DeLacy et al., 1996). Panter and Allen (1995) also concluded that BLUP was superior to least square means in predicting soybean cultivar performance, hence, in place of ordinary cell means, we used the BLUPs to draw biplots shown in this study. Hill and Rosenberger (1985) considered BLUP of genotype main effects in genotype-by-environment data and found BLUP to outperform BLUE. Yan et al. (2002) compared on-farm strip trials versus replicated performance trials with wheat (Triticum aestivum) based on BLUP of genotype effects. Yan and Rajcan (2003) used BLUP in soybean (Glycine max) based on a three-way model (genotype-by-location-by-year).

The cowpea breeding program of the International Institute of Tropical Agriculture (IITA) has been in the forefront leading in research and releasing varieties in Africa to provide high yielding and stable cowpea that targets the production area in SSA (Boukar et al., 2020). While GEI have been elaborately studied in cowpea to identify stable genotypes, no study exists to appraise and determine the suitability of test sites in selecting genotypes with superior and stable genotypes for grain yield in the traditional testing sites of IITA cowpea breeding program in Nigeria. The objectives of this study were to (i) evaluate variance components, heritability and estimate the importance and sensitivity of grain yield to GEI (ii) appraise the suitability of test sites used by IITA cowpea breeding for METs in terms of discriminatory and representative abilities and elucidate the nature of relationships existing between environments (iii) (elucidate/validate/assess the equivalence) and compare correlations among test sites as revealed by the BLUP-Based GGEBiplot, genotypic correlation and wards cluster analysis within the testing network.

## Materials and Methods

### MET Dataset and trial design

The cowpea breeding unit (CBU) of IITA routinely conducts Advance yield trials (AYTs), with each year having unique Entries. The datasets analyzed in this study were from the 2019 AYTs, which were clustered based on maturity into 6 Sets named Advance 1 to Advance 6 (Advance 1 being Extra Early, Advance 2 Early, advance 3, 4 and 5 medium and 6 late maturing/duration) Table 1. 32 datasets (i.e., 32 individual trials across the testing networks within the Advance sets) were assembled, curated and analysed for the study. Each Advance Set had 27 to 28 Entries including four checks (2 improved released lines and two local landrace) see **Table 1**, and tested in six sites (Advance sets 1,2,5 and 6) and four sites (Advance 3 and 5) Table 2. In Nigeria, the Program has four major testing sites; Minjibir, Kano, Shika and Ibadan (Table 2), two of these locations (Shika and Ibadan) can accommodate two cropping seasons within a year and therefore had two planting dates, such that a combination of location by planting dates constituted an environment (**Table 2**). Hence a maximum of 6 environments was used as a testing network. The coordinates and properties of each site/environment are described in Table 2. In each Environment, the Alpha lattice design was used to evaluate the entries. Because the 6 sets of Advances had three checks (See Table 1) in common, it was permissible to merge the sets into a Combined dataset for a Combined Advance analysis of variance to know if the Sets were unique.

**Table 1:** Advanced Yield trial and meta-data.

| Tabble 1: Advanced Yield trial and meta-data |  |  |  |  |  |  |  |  |
| --- | --- | --- | --- | --- | --- | --- | --- | --- |
| Trial | No. Entries | Design | Rep | No. Locations | Checks |  |  |  |
| Advance 1 (Set 1) | 28 | Alpha lattice | 3 | 6 | Achishiru | ALOKALOCAL | IT07K-297-13 | IT08K-150-12 |
| Advance 2 (Set 2) | 28 | Alpha lattice | 3 | 6 | Achishiru | ALOKALOCAL | IT07K-297-13 | IT08K-150-12 |
| Advance 3 (Set 3) | 28 | Alpha lattice | 3 | 4 | KANANNADO | DANILA | IT07K-297-13 | IT08K-150-12 |
| Advance 4 (Set 4) | 28 | Alpha lattice | 3 | 4 | KANANNADO | DANILA | IT07K-297-13 | IT08K-150-12 |
| Advance 5 (Set 5) | 28 | Alpha lattice | 3 | 6 | Achishiru | ALOKALOCAL | IT07K-297-13 | IT08K-150-12 |
| Advance 6 (Set 6) | 27 | Alpha lattice | 3 | 6 | Achishiru | ALOKALOCAL | IT07K-297-13 | IT08K-150-12 |

### Trial execution and Phenotypic data

**Table 1 also** shows the planting dates for each Advance set per location. Three to four seeds were planted and thinned to three stands at two weeks after emergence. Each plot consisted of four 4-meter rows. NPK fertilizers were applied at seedling stage across locations and the plots were sprayed with insecticides to control insect pests while hand weeding was done, when necessary, to control weeds. The two middle rows were considered during phenotyping, hence the land area for yield and other traits reported in this study was the two middle 4m length by 0.75 making a total land area of 6m^2^ (net plot). Plant stands at harvest (used as a covariate for yield) was determined by counting the number of stands in the net plot before harvest. At maturity, the net plots were harvested, threshed and weighed to obtain grain yield (GY) in grams per plot. The grain yield per plot was then converted to kilograms per hectare (kg/ha).

## Data analysis

### Distribution and correlation among traits

The R statistical software, version 3.5.2 (R Core Team 2018) was used to generate a graphical visualization using overlay-histograms and chats to show traits distribution and correlation between Advance sets. The means from each site were used to generate violin plots for individual sets to provide a graphical description of datasets in individual sites.

### 3.3 Means squares where computed in two stages

#### 3.3.1. Stage 1

Analysis of variance for combined Advance 2019 datasets: In order to determine if there were significant differences among the Advance sets, the Advances 1-6 across sites were combined, and since they both had 3 checks that were present in all datasets, it permitted ANOVA and estimation of error. The following model was fitted for the combined dataset, this was done for each trait:

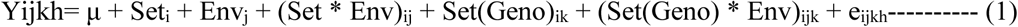

Where μ is the observed value of the ith genotype in the jth location, l is the general mean, Set_i_, Env_j_, (Set * Env)_ij_, Set(Geno)_ik_, (Set(Geno) * Env)_ijk_, and e_ijkh_ represent the effects of the genotype, location, the interaction etween genotype and location, the effect of genotypes nested within sets and the interaction between genotypes within set by location effect respectively.

#### 3.3.2. Stage 2

Analysis of variance for within Advance datasets: The second was to test for significance of main effects and Genotype by Environment interaction for each trait within each Advance set. The following models were implemented in R using agricolae and lme4 packages (Bates et al. 2015; Mendiburu 2020) to obtain mean squares (MS), coefficient of variations (CV) and standard errors of means for the trait. Both the Genotype and Environment were considered fixed effects and Significance of all effects was tested against mean square of error. Stand at harvest was used as a covariate for Grain Yield as shown in equation 2;

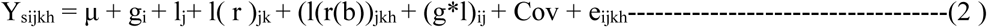

Where Y_sijkh_ is is the observed value of the ith genotype in the jth location, l is the general mean, gi, lj, l(r)_jk_, (l(r(b))_jkh_, (g*l)_ij_, COV, and e_ijkh_ represent the effects of the genotype, location, replication nested within location, block and replication nested within location, and the interaction between genotype and location, covariate and eijkh is the residual effect respectively. Since all Effects were fitted fixed, all effects were tested against the error term for significance.

### 3.4 Variance components and estimate of Broad sense Heritability

META R (Version 6.0) (Alvarado et al., 2015) was utilized to obtain estimates variance components for Genotype (G), Environment (E) and interaction between Genotype and Environment (GE) and Mean basis broad sense Heritability for all Within Advance datasets. Following model in equations 2 with Genotype fitted as random, META_R implements a linear mixed model in the LME4 R-package (Bates et al., 2015) that uses REML to estimate the variance components.

### 3.5. Phenotypic and genetic correlations among environments

Phenotypic and genetic correlations among environments within each Advance sets for grain yield was obtained using META-R (Alvarado et al., 2017) which uses the simple Pearson correlations to estimate the phenotypic correlation coefficients among environments and for genetic correlation coefficients it uses the equations from Cooper and Delacy (1994). We retained the default heritability threshold of 0.05.

### BLUP based GGEBiplot

Using the Lme4 library and Lmer function in R, the BLUPS for individual sites were obtained for grain yield using an Alpha-lattice design. We fitted the following model (equation Y) with Genotype as random effect; first for grain yield adjusted by a covariate

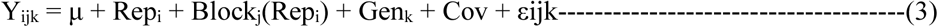

Where; Y_ijk_ is the trait of interest, μ is the overall mean effect, Rep_i_ is the effect of the ith replicate, Block_j_(Rep_i_) is the effect of the jth incomplete block within the ith replicate, Gen_k_ is the effect of the kth genotype, Cov is the effect of the covariate in the case of equation 3 for grain yield and εijk is the effect of the error associated with the ith replication, jth incomplete block, and kth genotype (Frutos et al., 2014). BLUPs obtained from equation 3 from each environment within each Advanceset where arranged in a two way table and using the GGEBiplotGUI package in R (Frutos et al., 2014) three biplots (which won where, Discriminativeness and representativeness and Relationship among environments) biplots relevant to explore the MET datasets were drawn (Yan et al., 2007). All biplots were not transformed (Transform=0), were environment centered (Centering =2) and were environment focused singular value partitioning (SVP=2). Except for the biplots Showing Relationship among Environments that was standard deviation-scaled (‘Scale=1’), all other biplots were not scaled (‘Scale=0’) (Yan and Tinker 2006, Yan and Holland 2010). These properties made the biplots suitable for appraising the appropriateness of the test environments and interrelationships among them (Yan and Tinker 2006).

The GGE biplots were used to delineate MEs based on the winning genotypes for each set (i.e., the polygons of extreme genotypes in the GGE). This strategy permits us to generate 6 biplots based on 6 different datasets which were taken to be different scenarios. The number of times a pair of environments shares a ME was used to define the historical ME structure. This strategy was used in lieu of year-to-year changes in ME patterns.

Following method described by Blanche and Meyers (2006), the mean distance of each site from the ideal location within each set were calculated and averaged across all six sets to obtain a ranking of environments based on idealness (discriminativeness and representativeness). Because the method is based on a standard error scaled biplot, it is very suitable to assess the idealness of the test sites (Yan and Holland 2010).

## Results

### 3.1 Trait distribution

Figure 1 shows trait distribution using an overlay histogram and correlation coefficients between Grain yeidl and other key agronomic traits within the 6 Advance sets. This was done to have a better understanding of the characteristics of grain yield and its relationship with other traits. Grain yield was generally skewed to the left for Advance sets 1-5 while a symmetric distribution was observed for Advance 6. A violin distribution within each environment is presented in supplementary material 2 and it showed that variable trait distribution existed within each Advanced sets for individual location.

**Figure 1:**
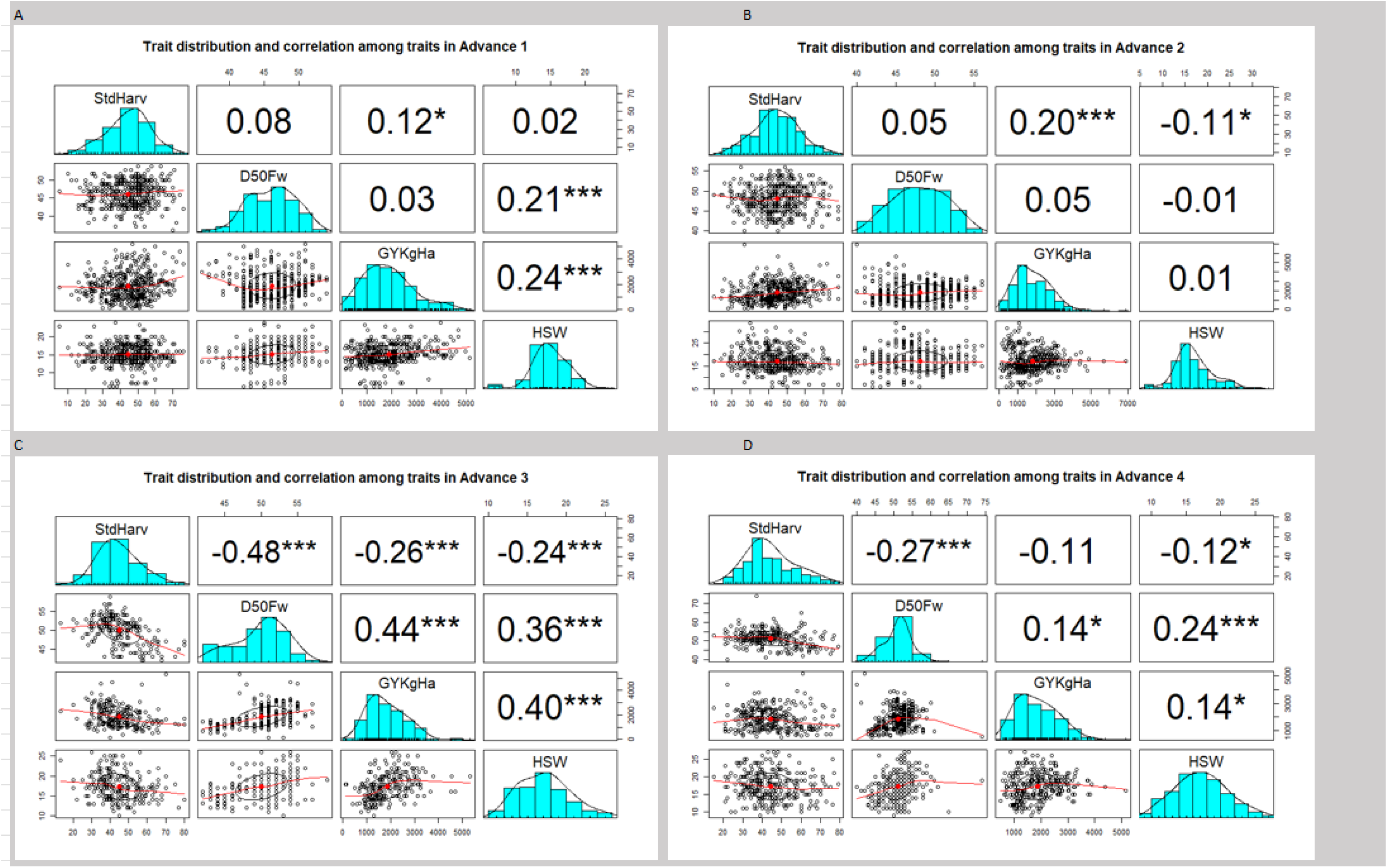
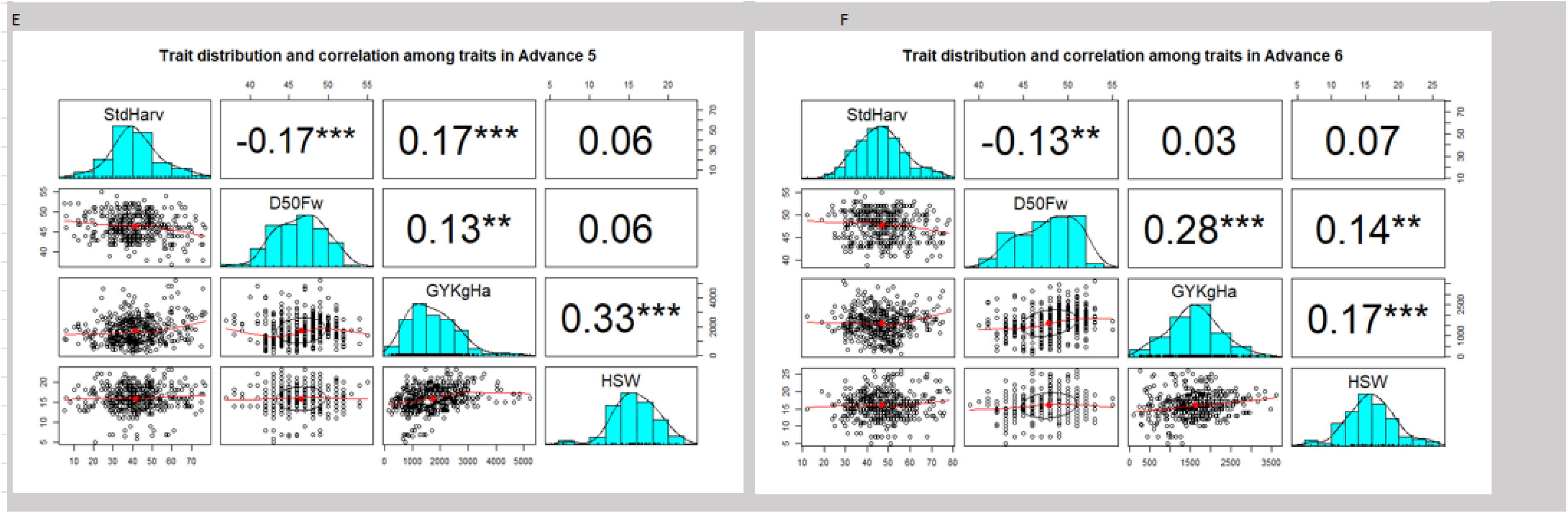
Trait distribution and correlation.

Significant weak positive correlation was observed between Stand at harvest and grain yield in Advance sets 1 (r=0.12, P<0.05), Set2 (r=0.2, P<0.01) and Set 5 (r=0.17, P<0.001), but a significant weak negative correlation was observed in Adv 3 (r=-0.26, P<0.001). The correlation between grain yield other traits exempting stand at harvest was consistently weak and positive. Similarly, the relationship was observed between Grain yield and D50FW was consistently weak and positive (SIGBNIFICANT OR NOT state pleses).

### Means squares and coefficients of variability

Between Sets Combined Analysis of variance: Table 3 shows the combined ANOVA. The combined analysis of variance to test for differences between Sets revealed that the Sets were significantly (P<0.001). It also showed that the Interaction between Genotypes and Sets was significant (P<0.05).

**Table 3:**
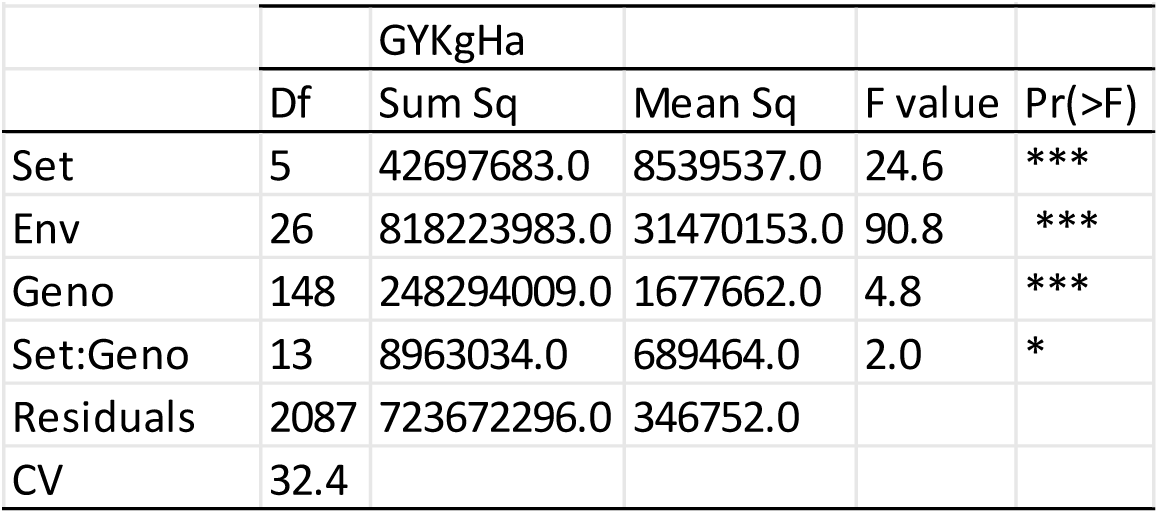
Combined-sets ANOVA (Between Sets)

#### Within Advance Sets

Table 4 shows the analysis of variance within the Sets. For grain yield, the GE interaction effects were highly significant (P<0.001) in Advance sets 1-5 and significant at P<0.01 for Advance 6. CV ranged from 23.7% (Adv 3) to 29.9% (Adv 2).

**Table 4:**
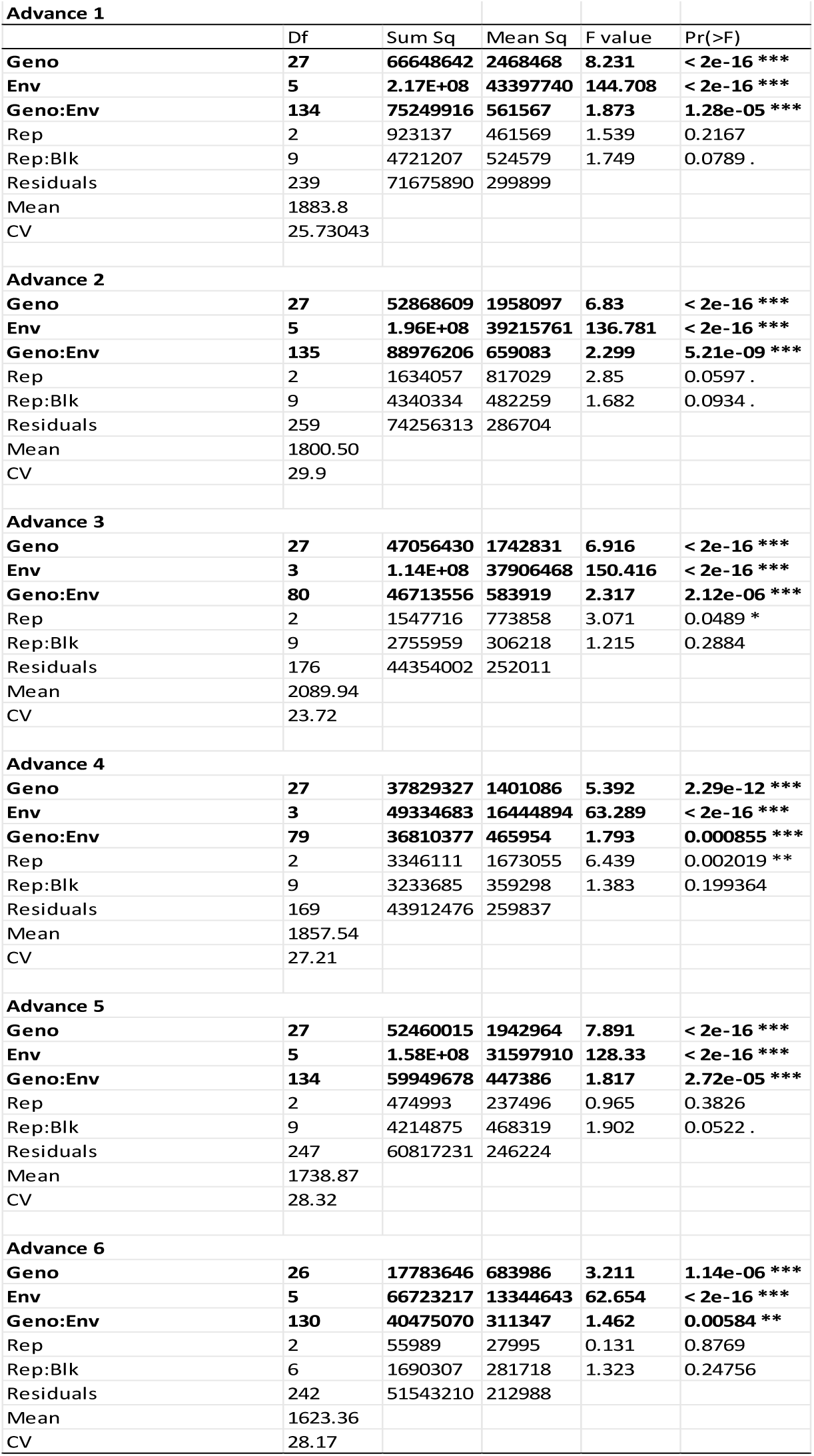
ANOVA within Sets.

### Variance component and heritability

Table 5 shows the variance components, percent contribution and heritability estimates. The variance components were expressed in percentage to provide a more succinct picture of what proportion each main and interaction effect was explaining of the total phenotypic observation. Environment accounted for most of the variation observed ranging from 58% (Adv4) to 73.9% (Adv3), followed by GE interaction with the range of 16.1% (Adv 5) to 22.6% (Adv4). G had the least proportion ranging from 9.5% (Adv6) to 19.3% (Adv4), hence, GE explained more of the phenotypic variation captured than G; with Advance 4 having the highest ratio of G (19.3%) to GE (22.6%). Broad sense heritability for grain yield were moderate to high with a range of 0.53 (Adv 6) to 0.7 (Adv 1 and 5).

**Table 5:**
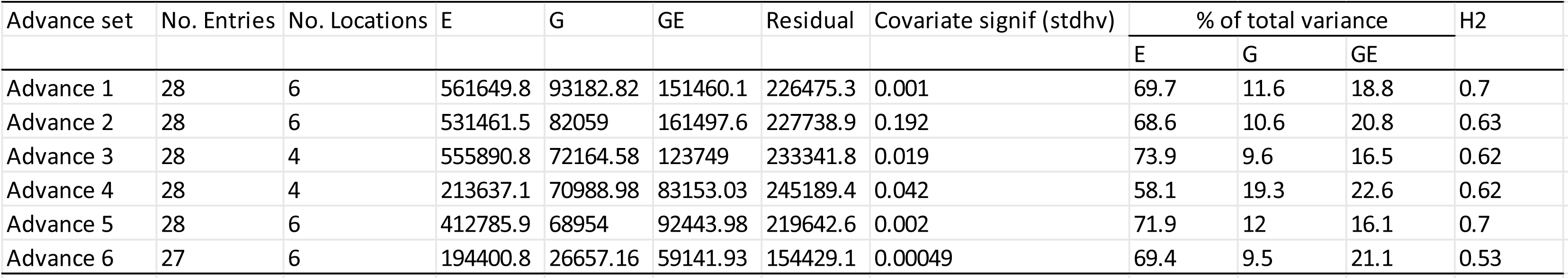
Variance component and broad sense heritability.

### Genetic and phenotypic correlation among environments

Table 6 shows the Phenotypic (left) and Genetic (right) correlations among environments for Grain yield.

**Table 6:** Correlations. correlation among environments for Grain Yield

| Phenotypic correlation among environments |  |  |  |  |  |  | Genetic correlation among environments |  |  |  |  |  |  |
| --- | --- | --- | --- | --- | --- | --- | --- | --- | --- | --- | --- | --- | --- |
| Adv1 | Env | E2 | E6 | E3 | E4 | E1 | Adv1 | Env | E2 | E6 | E3 | E4 | E1 |
|  | E6 | -0.09 |  |  |  |  |  | E6 | -0.17 |  |  |  |  |
|  | E3 | 0.13 | 0.46* |  |  |  |  | E3 | 0.20 | 0.92** |  |  |  |
|  | E4 | 0.36 | 0.27 | 0.47* |  |  |  | E4 | 0.47 | 0.47 | 0.66 |  |  |
|  | E1 | 0.08 | 0.13 | 0.53** | 0.58** |  |  | E1 | 0.10 | 0.22 | 0.75 | 0.70 |  |
|  | E5 | 0.40* | 0.42* | 0.34 | 0.47* | 0.46* |  | E5 | 0.51 | 0.69 | 0.47 | 0.56 | 0.52 |
| Adv2 | Env | E2 | E6 | E3 | E4 | E1 | Adv2 | Env | E2 | E6 | E3 | E4 | E1 |
|  | E6 | 0.15 |  |  |  |  |  | E6 | 0.25 |  |  |  |  |
|  | E3 | 0.19 | -0.31 |  |  |  |  | E3 | NA | NA |  |  |  |
|  | E4 | 0.26 | 0.34 | 0.07 |  |  |  | E4 | 0.44 | 0.44 | NA |  |  |
|  | E1 | 0.01 | 0.28 | -0.01 | 0.19 |  |  | E1 | 0.01 | 0.35 | NA | 0.24 |  |
|  | E5 | 0.17 | -0.11 | 0.32 | 0.35 | 0.69*** |  | E5 | 0.26 | -0.13 | NA | 0.43 | 0.80 |
| Adv3 | Env | E1 | E4 | E2 |  |  | Adv3 | Env | E1 | E4 | E2 |  |  |
|  | E4 | 0.30 |  |  |  |  |  | E4 | 0.38 |  |  |  |  |
|  | E2 | -0.22 | 0.16 |  |  |  |  | E2 | -0.30 | 0.23 |  |  |  |
|  | E3 | 0.67*** | 0.38* | 0.05 |  |  |  | E3 | 0.77 | 0.48 | 0.07 |  |  |
| Adv4 | Env | E1 | E4 | E2 |  |  | Adv4 | Env | E1 | E4 | E2 |  |  |
|  | E4 | 0.11 |  |  |  |  |  | E4 | 0.15 |  |  |  |  |
|  | E2 | 0.03 | 0.45* |  |  |  |  | E2 | 0.05 | 0.71 |  |  |  |
|  | E3 | 0.21 | 0.39* | 0.06 |  |  |  | E3 | 0.45 | 0.88 | 0.14 |  |  |
| Adv5 | Env | E5 | E1 | E4 | E2 | E6 | Adv5 | Env | E5 | E1 | E4 | E2 | E6 |
|  | E1 | 0.65*** |  |  |  |  |  | E1 | 0.78 |  |  |  |  |
|  | E4 | 0.24 | 0.52** |  |  |  |  | E4 | 0.29 | 0.61 |  |  |  |
|  | E2 | 0.42* | 0.11 | 0.00 |  |  |  | E2 | 0.90* | 0.23 | 0.01 |  |  |
|  | E6 | 0.26 | 0.31 | 0.14 | -0.21 |  |  | E6 | 0.42 | 0.48 | 0.22 | -0.59 |  |
|  | E3 | 0.35 | 0.57** | 0.53** | -0.02 | 0.17 |  | E3 | 0.43 | 0.68 | 0.66 | -0.03 | 0.27 |
| Adv6 | Env | E5 | E1 | E4 | E2 | E6 | Adv6 | Env | E5 | E1 | E4 | E2 | E6 |
|  | E1 | 0.44* |  |  |  |  |  | E1 | 0.59 |  |  |  |  |
|  | E4 | 0.15 | 0.34 |  |  |  |  | E4 | NA | NA |  |  |  |
|  | E2 | -0.28 | -0.09 | -0.44* |  |  |  | E2 | -0.92** | -0.33 | NA |  |  |
|  | E6 | 0.13 | 0.22 | 0.29 | 0.16 |  |  | E6 | 0.18 | 0.31 | NA | 0.58 |  |
|  | E3 | 0.23 | 0.42 | 0.51** | -0.36 | 0.23 |  | E3 | 0.30 | 0.56 | NA | -1.00*** | 0.33 |

### Genetic correlation

Adv1 indicated a significant (P>.001) strong positive genetic correlation between (r=.92) between E3 and E6. No significant r values were observed for Adv 2, 3 and 4 for genetic correlation. Adv 5 however showed a significant (P>.05) strong positive genetic correlation between (r=.9) between E2 and E5. Adv 6 was interesting as it showed that E2 had a strong negative significant correlation E5 (r=-.92) and E3 (r=.99) contradicting Adv 5.

### Phenotypic correlation

The correlation coefficient was ranged from weak to strong, except for Adv6 (r=-0.44) E4 and E2. E1 and E5 had a consistently significant r values ranging from moderate (Adv 6 r=0.44) to strong (Adv 2 r=0.69), while E3 and E6 were mostly not significant at P<0.05 and variable; weak for Adv1 (r=0.46) and negative in Adv 2 (r=-0.31 NS). E4 and E2 had a consistent weak relationship that was negative and significant Adv 6 (-0.44) but positive in Adv 4 (0.45) and zero but not significant relationship was observed for Adv5.

Evaluation of Test Environments: Table 7, shows the summary of number of sectors and the environments forming each sector for Adv1-6 for grain yield. Biplots are presented in supplementary material 2.

**Table 7:** Summary of ‘which won where’ and Mega environments. Synthesis of Sectors delinating Mega environemnts for Grain yield

| Grain yield | Sector 1 | Sector 2 | Sector 3 |
| --- | --- | --- | --- |
| Advance Set 1 | E1, E2, E3, E4, E5 and E6 |  |  |
| Advance Set 2 | E2, E3, E4 and E6 | E1 and E5 |  |
| Advance Set 3 | E2 | E4 and E3 | E1 |
| Advance Set 4 | E1 | E2, E3, and E4 |  |
| Advance Set 5 | E4 | E1, E2, E3 and E6 | E5 |
| Advance Set 6 | E2, E3, E4 and E6 | E1 | E5 |

In Adv 1 all the environments fell into a single sector. Adv 2 had two sectors; Sector one with E2, E3, E4 and E6 while E4 and E5 formed the second. Adv 3 had 3 sectors with environment markers in them; sector 1 had E2 alone, sector 2 had E4 and E3 while E1 alone was placed in the third sector. Adv 4 had 2 sectors; E1 alone in the first while E2, E3 and E4 formed the second. Adv5 had three sectors E4 alone formed a sector, E1, E2, E3, and E6 formed the second while E5 formed the third. Finally, Adv 6 which had 3 sectors with E2, E3, E4 and E6 forming a sector, while E1 and E5 each formed its own sector. While E1 and E5 fell into different sectors in Adv5 and 6, E3 and E6 were consistently in the same sector. Sets 1, 2, 3, 4 and 5 had IT08K-150-12 a check as the vertex genotype in all the Mega environments where E4 was found; the implication of this is further elaborated in the discussion section.

Figure 2 shows the discriminatory and representativeness of test environments for Grain yield for sets evaluated. The discriminatory and representative ability of each environment within each Sets are summarized in Table 8 with biplots shown in supplementary material 2. With respect to discriminatory ability, E1 showed moderate (Adv 4,5 and 6) to high (Adv 2 and 3) discriminatory ability for grain yield, E2 showed high discriminatory ability in Set 1,3 and moderate in set 4 but poor in sets 2, 5 and 6. E3 and E6 were generally poor in terms of discriminating among genotype performance having a fair performance only Adv 6. E4 was the most discriminating environment, although it had a very short vector in Adv 6. E5 showed a moderate to high discriminative ability.

**Figure 2:**
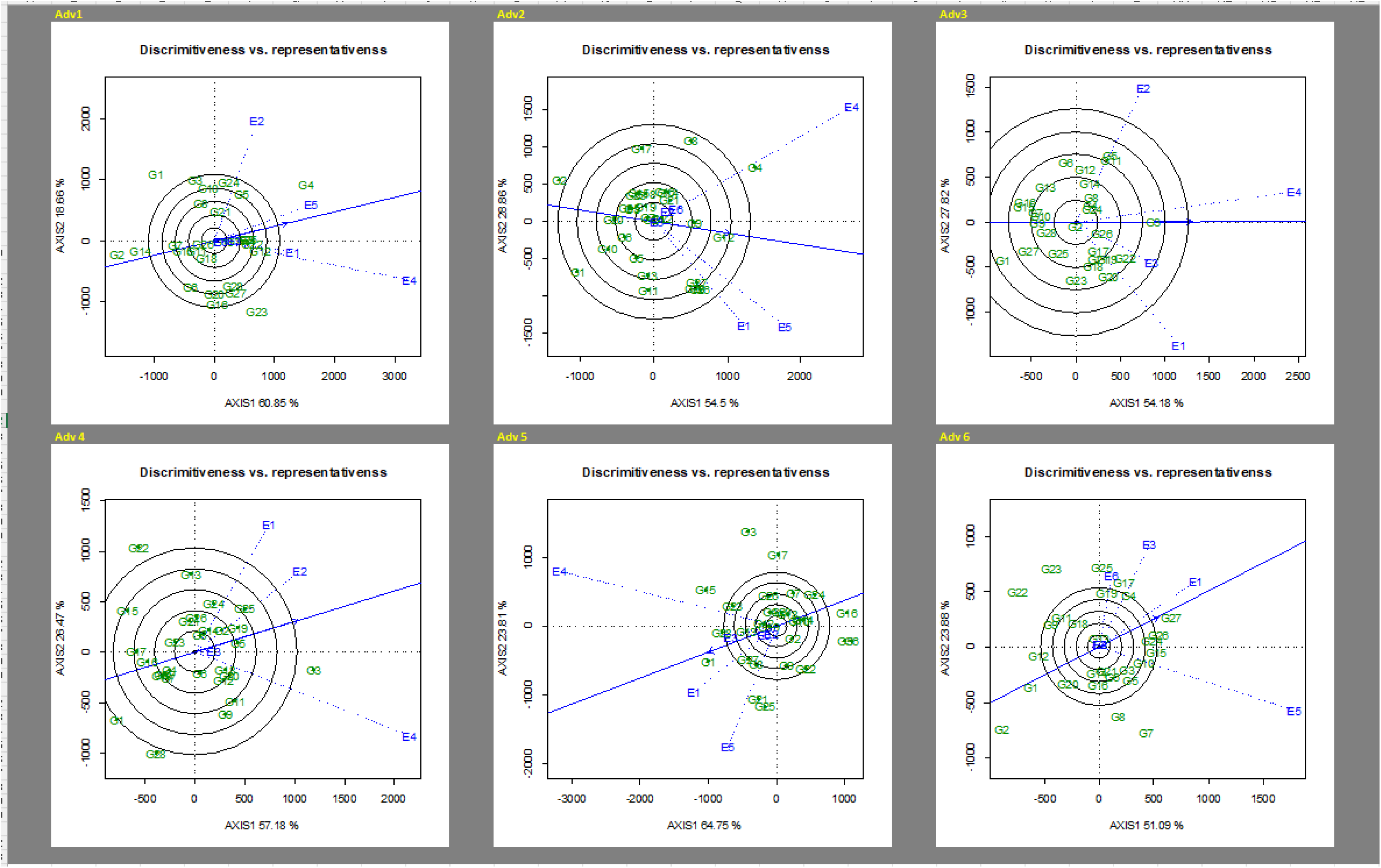
Discriminatory and representative ability of Environments.

**Table 8:** Summary of Discriminative and representativeness of test locations.

| Environment | Set 1 | Set 2 | Set 3 | Set 4 | Set 5 | Set 6 |
| --- | --- | --- | --- | --- | --- | --- |
| E1 | PD;MR | HD;PR | HD;PR | MD;PR | MD;HR | MD;HR |
| E2 | HD;PR | PD;PR | HD;PR | MD;MR | PD;PR | PD;PR |
| E3 | PD;PR | PD;PR | PD;MR | PD;PR | PD;HR | MD;MR |
| E4 | HD;MR | HD;PR | HD;HR | HD;PR | HD;PR | PD |
| E5 | MD;HR | HD;MR |  |  | MD;PR | HD;PR |
| E6 | PD;PR | PD;PR |  |  | PD;PR | PD;PR |

In terms of representativeness, for grain yield, E1 showed desirable level of representativeness only is Set 1, 5 and 6. E2 showed suitable level of representativeness only in Advance 4, due to its very short vector in other plots. E3 showed a high level of representativeness in set 5 and moderate in sets 3 and 6. E4 was consistently moderate in all the biplots except in Set 3 where it showed a good. E5 showed a desirable level for representativeness in Sets 1 and 5 but less desirable in Sets 5 and 6. E6 conxsistently showed a short vector and was therefore not desirable for grain yield assessment.

### Relationship among Environments

The relationship among environments was determined by the size of the angle between the vectors of any two environments. The larger the angle, the less correlated the environments. If the angle is acute (\90_), it indicates a strong positive correlation between the environments, suggesting that the same information about the genotypes could be obtained from correlated test environments without sacrificing precision. If the angle is a right angle (=90_), no relationship was indicated, if the angle was obtuse ([90_), it indicated a strong negative correlation and an indication of the presence of a strong crossover GE, and if the angle was on a straight line (=180_), it indicated a no relationship/correlation (Yan and Tinker 2006).

Figure 3, shows the relationship among Environments for grain yield.

**Figure 3:**
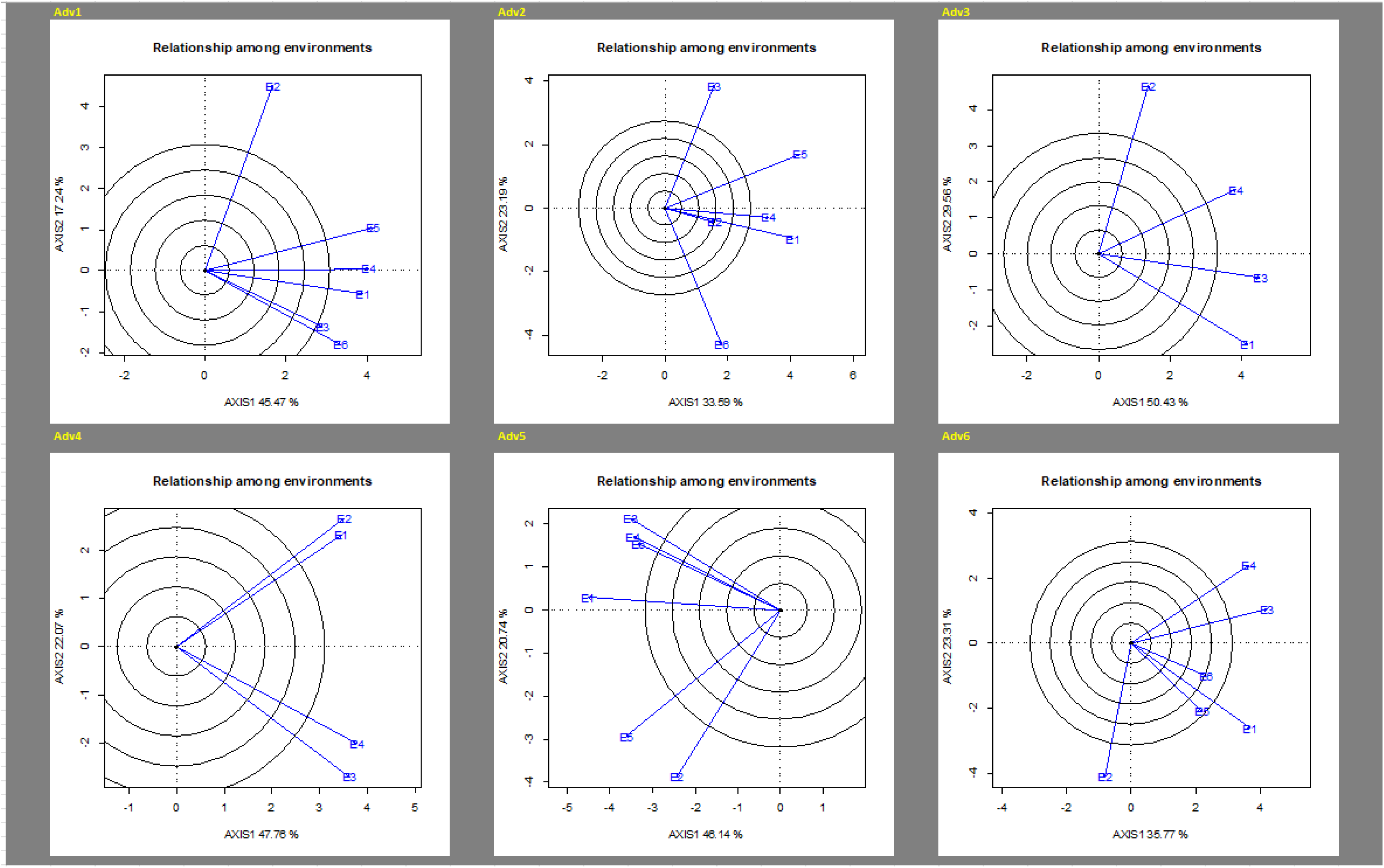
Relationship among Environments.

**Figure 4:**
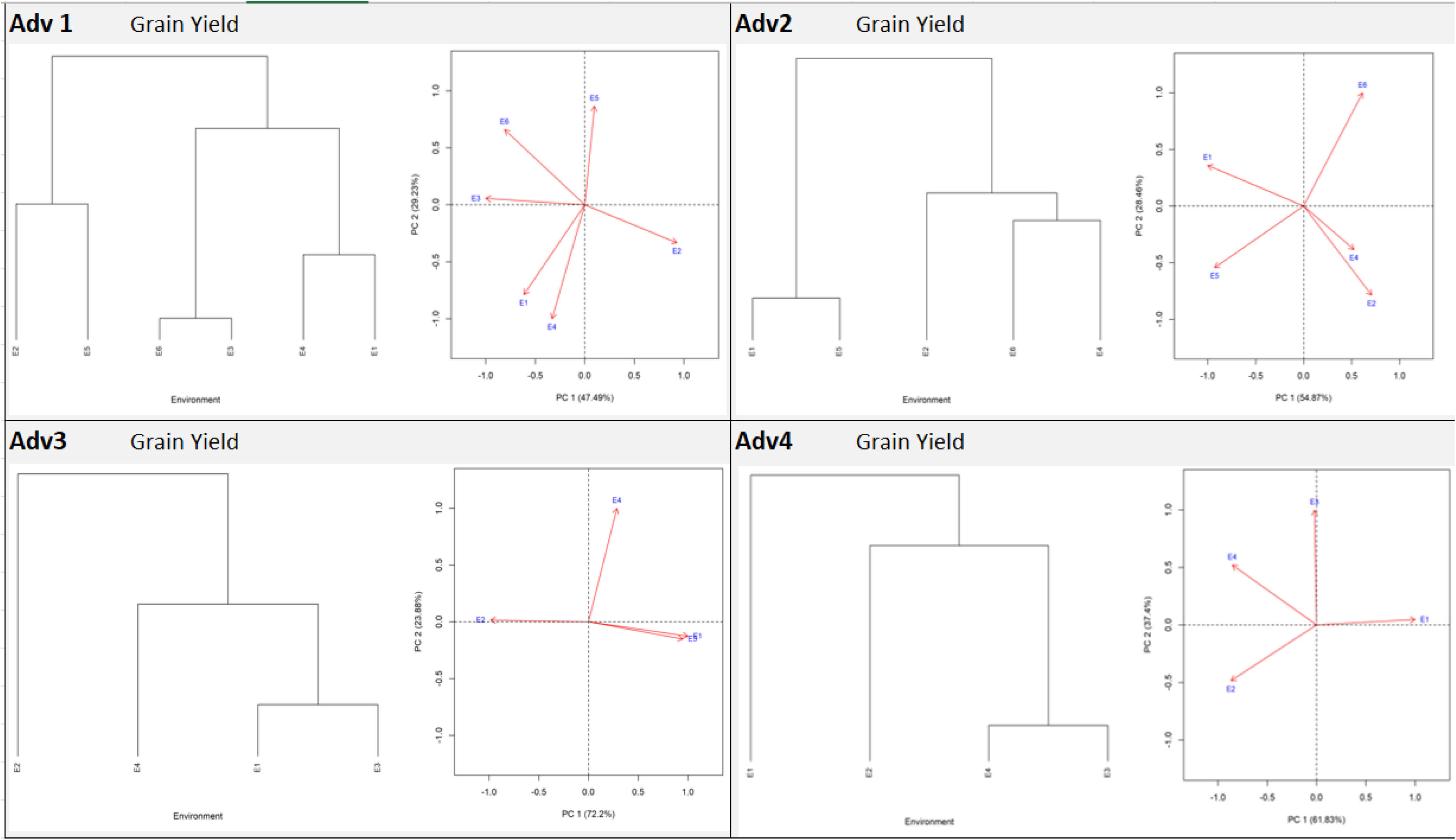
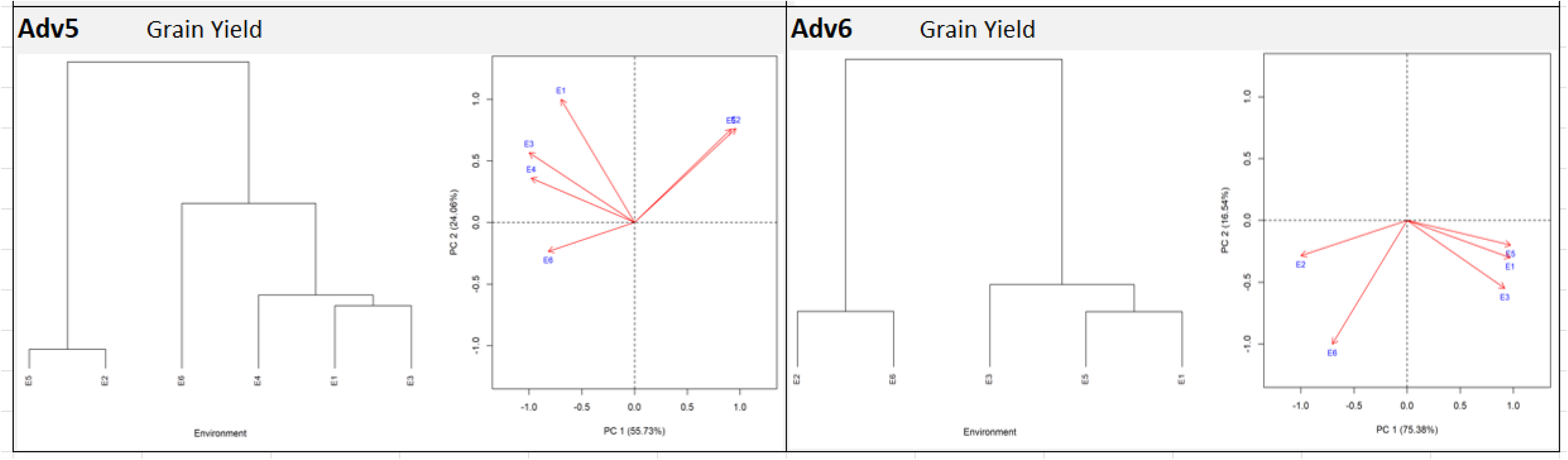
Dendrogram.

**Figure 5:**
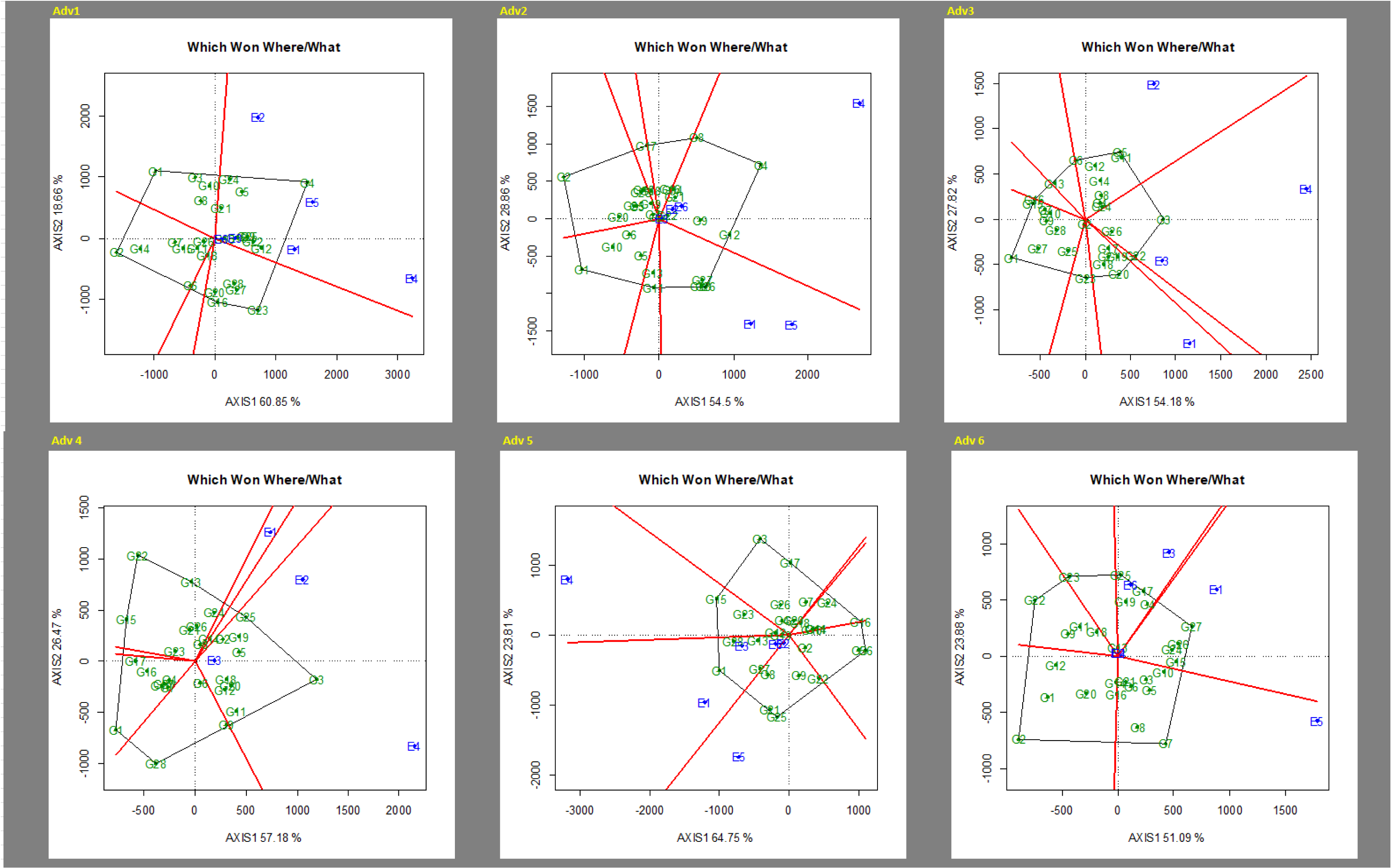
Mega Environment.

**Figure 6:**
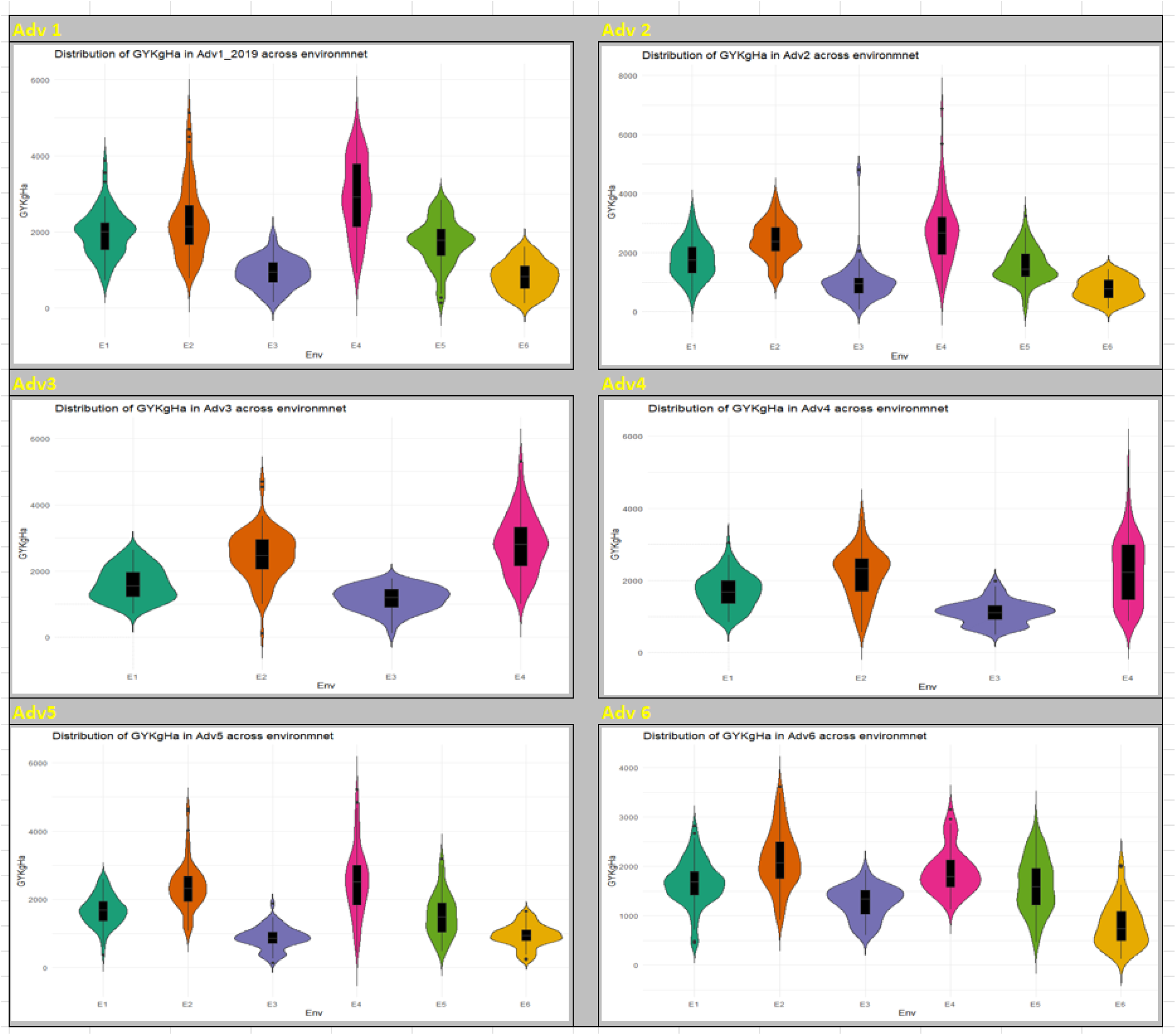
Distribution of traits.

Adv1 E2 was negatively correlated with E3 and E6, while its relationship with other sites was positive but weak. E1, E3, E4, E5 and E6 were all closely related to one another, with E3 and E6 having the closest relationship. Adv2: E3 and E5 were both negatively correlated with E6. On the other hand, E1, E2 and E4 were positive and strongly related. E5 had a weak positive relationship with E1 and E3. Adv3: A negative relationship was observed between E2 and E1, while a weak positive relationship was observed between E2 and E3. E1, E3 and E4 all had a positive relationship within themselves. Adv4: showed two distinct groups with a weak positive association between the two groups. In the first group, E1 and E2 were closely related while E3 and E4 were also closely related on the other hand. Adv5: E3 and E2 showed a negative association as observed in Adv1 and 2. E2 also had a strong positive association with E5 but weak positive association with E1, 4 and 6. Adv6: E2 had a negative association with all other sites. E4 and E5 shows a very weak association, while E1, 5 and 6 showed a strong positive association among themselves.

Table 9 shows the suitability of the test sites averaged across the six Advance sets following method described by Blanche and Myers 2006. The standardize distance between actual and ideal location were averaged and ranked. The smaller the distance, the more ideal the location is hence, Shika August with the lowest average distance of 27.02 and standard deviation (SD) of 5.75 ranked first. It was closely followed by Shika June which ranked second; with an average distance of 28.09 with SD of 8.04. Ibadan May was with the highest distance ranked last with an average distance of 42.96 and SD of 5.06

**Table 9:**
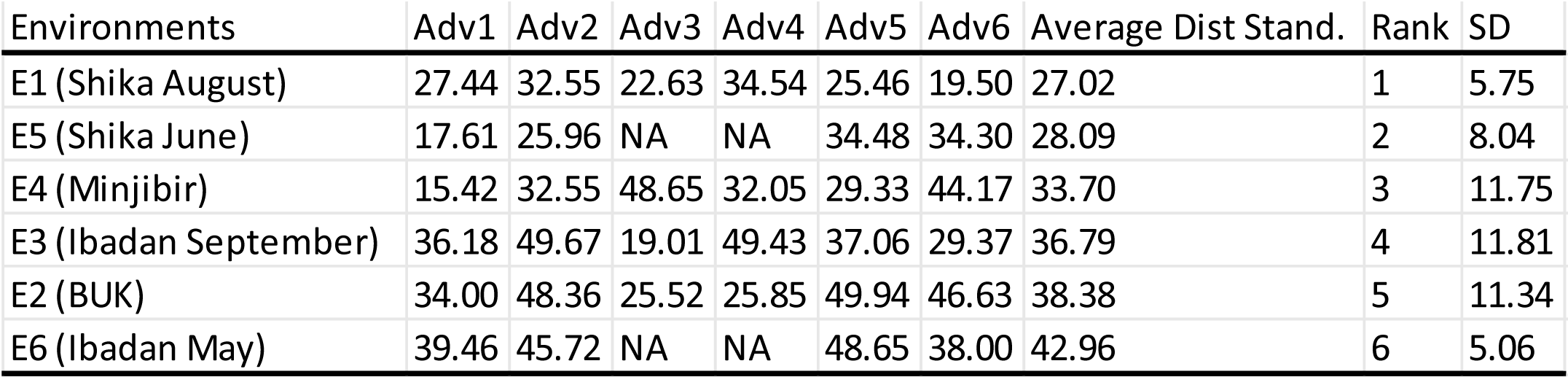
Rankiing of Environments based on closeness to the ideal site.

### Dendrogram cluster analysis

GY Cluster plots: Adv1 showed that all the environments fell into 3 clusters; E1 and E4, E3 and E6 and E2 and E5 with stronger similarity with each other within each cluster. Adv2 showed that E1 and E5 are similar while E4 and E6 were clustered together with E2, with E4 and E6 been more similar. Adv3 showed that E2 was unique as it formed a leaf alone, while E4, E1 and E3 were clustered and are more similar, with E3 and E1 been more closely related. Adv 4 showed E1 to be unique while E2,3 and 4 were clustered together. In Adv5 E5 and E2 were clustered together while other sites formed another cluster in which E1 and E3 where more related to each other than E4 and E6 which was in a unique leaf. Adv6 reveal a close relationship between E2 and E6, while E1 and E5 along with E3 (furthest) formed a cluster.

Equivalence with GGE biplot: Comparing results from the different methods we used to evaluate relationship and clustering among environments reveals that there is a perfect similarity in the direction of correlations for grain yield between genetic correlation and GGE biplot; as positive relationships reported by genetic correlation were also positive according to the GGE biplots for grain yield.

In terms of clustering of environments in terms of similarity; the Ward’s cluster Dendrogram and GGE biplot had a perfect synergy in Adv 3 and 5 to an appreciable extent; as the clustering by GGE biplot happens to be exact with the Ward’s method for grain yield.

## Discussion of results

The representations from overlay histogram showed a continuous distribution for all GY corroborative of quantitative traits and indicates the presence of multiple genes are in play with strong environmental influence (Ongom et al, 2021, Chuan et al., 2009, Kelker et al., 1986 and Sinnott 1937) and this agrees with the opinion of Roy (2000) and Jayaramachandran et al (2010) which say that quantitative characters in plants whose spread extends to the left or right show the influence of the environment, genotype and environment interaction. Therefore, in order to maximize the genetic gain in respect of these traits with positively skewed distribution there will be need for intense selection from the existing variability (Roy, 2000). All traits were symmetric except for Grain yield that was skewed to the right in Advance Sets 1-5; signifying that there were few Entries that had higher values than the mean, mode and median, it was confirmed through careful curation that these values are not outliers because the Entries had a consistent high grain yield across the three reps. Therefore, this high yielding Entries are potential lines to concentrate on when assessing the stability of the Entries (which is not the focus of the study). A violin plots showed that there were different patterns of response within each Set across the locations. The distributions observed in this study were similar to a recent study (Ongom et al., 2021) and other prior studies (Boukar et al., 2016).

A major constraint in the effort to identify superior cowpea lines with narrow or wide adaptation is the disparity in response of the tested genotypes to the various environmental conditions in which the plants are grown (Badu-Apraku et al., 2013), hence the significant mean squares obtained from our results in the present study for genotype main effect signified differential responses of the genotypes to environments necessitating the need to identify high-yielding and stable genotypes across the test environments (Badu-Apraku et al. 2013, Moghaddam and Pourdad 2009). The presence of a highly significant GEI across the maturity groups is a confirmation of the need for extensive testing of cultivars in multiple environments before they are being recommended for release. This also affirms the need for cowpea breeders to take GEI into serious consideration when evaluating potential lines and to always pay attention to its magnitude, relative to the magnitude of G and E effects for grain yield. Environment main effect were highly significant indicating uniqueness of the environments in terms of differences in magnitude of biotic and abiotic factors within the growing period. Dissection of the total phenotypic variance into main and interaction components showed that the environmental variance accounted for 58.1% - 73.9% of the variation for grain yield with the genotype contributing only 9.5% - 19.3%, thus, reflecting a much wider range of environmental main effects over genotypic main effects. This finding is in agreement with the results of most multi-environment trials, (DeLacy et al. 1996; Cooper et al. 1994; Badu-Apraku et al. 2011). In all cases GxE variance component was higher than Genotypic Variance, signifying that genetic variability was masked by higher GxE interaction. This made the interpretation of the main effects not meaningful considering the Statistical marginality condition (Girma and Makumbi, 2014).

By combining the Advances sets into a single dataset, we were able to conduct an ANOVA which shows that the sets were statistically different, validating our approach to use datasets from multiple sets to address the objectives of the study. In addition, it also shows that grouping of the Entries based on Maturity group was efficient in categorizing the entries. This further confirms the submission of Yan and Rajcan 2003 that single-year multi location datasets are sufficient in genotype evaluation as multi-year datasets. Cross and Helm 1986, Gellner 1989, and Bowman, 1998 all concluded that it was sufficient to use single-year multiple location data sets for selection.

According to Manggoel et al. (2012) and Rashwan (2010), high broad sense heritability values usually indicate the predominance of additive gene action in the expression of the traits. This further supports the practice of using single seed decent to improve GY in cowpea. The high values for broad-sense heritability will help in transferring the genetic characteristics from the parents to offspring (Rashwan 2010).

Mega environments: Our results suggest that about 3 possible mega environments exist for GY. However, this is not sacrosanct as many years of repeated combination of treatments are required to establish a ME (Yan and Holland 2010). However, it is worth mentioning that for grain yield which is the main trait for selecting a line, Ibadan May and Ibadan September consistently fell in the same ME for all six biplots drawn supporting one of our main conclusions that one of them is a redundant environment that could be dropped in order to optimize the MET network and as a cost saving strategy. Shika June and Shika August fell into different MEs for late and medium maturity groups but formed a ME together for the early maturity group. This suggest that Shika June and August can be retained for medium and late maturity groups while one of them could be dropped for the early testing groups in an optimized MET network.

Discrimination and representativeness: The ability of a test site to maximize the variance among genotypes is referred to discriminating ability, hence a good test site should have a high level of discriminating ability (Blanche and Myers 2006). Using the GGE biplot, it is measured by approximating the length of the site vectors and with the average environment axis (AEA) (Yan and Tinker 2006) the level of representativeness of the test sites can be determined; a test site that has a smaller angle with the AEA is more representative of the other test sites (Yan and Tinker 2006). Hence for a site to be ideal for MET it should be both discriminating and representative. Discriminating but non-representative test environments are not useful for selecting broadly adapted genotypes and will only lead to a poor recommendation of varieties, they can only be useful for selecting specifically adapted varieties if the target environments can be divided into mega-environments” (Yan and Tinker 2006).For grain yield, it was obvious that Ibadan May which consistently showed very poor discriminative ability for grain yield does not meet the requirements for inclusion in a MET network. While Minjibir was consistently discriminative and the most discriminative environment in this study, it had a poor representation and it implies that it is only fit for selecting specifically adapted varieties if it is identified as a mega environment or culling of unstable genotypes (Yan et al 2007). Badu-Apraku et al., 2013 reported similar instances where an environment was highly discriminative but had poor representativeness in maize. Omogui et al 2017 also showed that Minjibir was highly discriminative but a had fair to poor representativeness however they concluded that Minjibir was a good site but not good enough to be identified as an ideal site in their study. Ibadan September had poor discriminative ability when evaluated for the early maturity groups but improved and showed moderate discriminativeness for medium and late maturity groups implying that it may be useful for testing medium and late duration lines. Shika June and August showed good discriminativeness across Sets (and maturity groups) by having a moderate vector length. BUK was inconsistent in its discriminative ability; it showed good discriminativeness in Sets1,3 and 4 but poor in Sets 2,5 and 6.

Idealness of test sites: The Blanche method (Blanche and Mayers 2006) is an effective statistic to rank tests sites based on the two main criteria; discriminativeness and representativeness and it has been used by researchers to rank environments for instance, Yan et al., 2010, used it to rank environments used in soybean production in southern Wisconsin, Xu et al., 2014 used same method to appraise the idealness of test sites used in cotton production in the Yangtze River Valley (YaRV), China. Also according to Yan and Holland 2010, conclusions from the this method are very relevant and well related to the theory of indirect selection. From our results, Shika August and June ranked first and second respectively with a low Sd indicating that the low distances between the sites and ideal location had little variance. Minjibir was ranked third with the second highest SD of 11.75 indicating that it was not stable in terms of ranking across the six sets, an extreme example of its inconsistently is seen in Adv 1 where it ranked best and Adv 3 where it was ranked worst. BUK had a similar trend with Minjibir and had the highest SD showing its inconsistency and it was ranked second to the worst. Ibadan May was ranked last indicating that it was least desirable. It also had a low SD indicating that it consistently ranked poorly across the six sets and should be removed from the testing network. Ibadan September on the other hand ranked fourth with a high SD also indicating that it was not consistent in its ranking across the six sets; ranking worst in Adv 2 and 4. Ibadan September could be dropped too due to its inconsistency and poor discriminating ability.

### Relationship among environment

In this study, the relationship among environments was studied via the biplot and correlation coefficients. High genetic correlation among Environment suggests little GE effect among Test sites (Yan and Tinker 2006). Grain yield showed higher impact of GEI and hence recorded lower correlation coefficients. Also, a High genetic correlation between two environments implies that genotypes had similar ranks in the two, hence the two locations provide similar information regarding variety performance. it implies a similar ranking of the genotypes in these pairs of locations, hence Ibadan May and Ibadan September that were consistently positively correlated implies that one of the locations could be dropped in this case Ibadan May which had the poorest discriminative ability.

### Equivalence among GGE biplot, correlation cluster analysis

Brankovic et al., 2012, reported disagreement between the relationships among test sites revealed by biplot did not always coincide with Spearman’s rank correlation coefficients for site pairs. Our study had a contrasting results. A perfect congruent was seen between the results from the biplot and correlation as well as cluster analysis. For instance, Significant relationship between E3 and E6 was strong in both biplot and correlation coefficients for Adv1 and negative relation between E2 and E6 was observed in both biplot and correlation. Abe et al., 2015 used cluster analysis and reported its efficiency in studying relationships which is a similar trend observed in our study.

## Conclusion

The results of this study contribute to the body of literature on MET analysis in cowpea that indicates that GEI is important and plays a huge role in yield. This is the first study that looked into details suitability of the IITA cowpea breeding program METs network based on their discriminative and representativeness. Our study identified Ibadan May as redundant while Minjibir was highly discriminative was but not representative. Gy was also confirmed to be very sensitive to GEI. A strong equivalence was discovered between GGE biplot and Genotypic correlation and cluster analysis, indicating that any of the methods could be used in describing relationship among environments. Ibadan May was consistently found to be poorly discriminating and consistently paired with Ibadan September suggesting that it is a redundant environment and consistently formed a ME together and should be dropped. Shika June and Shika August were found to be unique and highly discriminating environments and should be retained. Therefore, there is a need to sample more testing sites using similar principles applied in this study to identify sites with high representativeness, discriminating ability, and repeatability for use in evaluating and selecting superior cowpea genotypes.

Because both static and dynamic factors determine the discriminativeness of an environment. In the current study no such data was considered. Future studies should focus on including some if not all of these factors, this will provide a detail cause of GEI in cowpea METS and extend better informed options on how to manage GEI in cowpea breeding programs. Multiple trait selection is crucial to variety adoption. The current study looked at only grain yield, its well known that breeders don’t release a variety based on a single trait nor do farmers adopt a variety based on a single trait, hence we recommend further studies which considers a traits index selection stability as suggested in Falcon et al., 2020. Finally on limitation in our study, we used Advanced trials which contains genotypes that are highly fixed. A study with more genetically variable population which maximizes the variability and has a larger range is recommended. Finally, because multiple years are required to confirm the existence of mega environments, future studies with year effect should be considered.

## Notes

### Competing Interest Statement

The authors have declared no competing interest.

### Funding

- IITA Cowpea Breeding Unit, via AVISA Project

